# Constitutive expression of the Type VI secretion system carries no measurable fitness cost in *Vibrio cholerae*

**DOI:** 10.1101/2023.03.24.534098

**Authors:** Christopher Zhang, William C. Ratcliff, Brian K. Hammer

## Abstract

The Type VI Secretion System (T6SS) is a widespread and highly effective mechanism of microbial warfare; it confers the ability to efficiently kill susceptible cells within close proximity. Due to its large physical size, complexity, and ballistic basis for intoxication it has widely been assumed to incur significant growth costs in the absence of improved competitive outcomes. In this study, we precisely examine the fitness costs of constitutive T6SS firing in the bacterium *Vibrio cholerae*. We find that, contrary to expectations, constitutive use of T6SS has a negligible impact on growth, reducing growth fitness by 0.025 ± 0.5% (95% CI) relative to a T6SS-control. Mathematical modeling of microbial populations demonstrates that, due to clonal interference, constitutive expression of the T6SS will often be neutral, with little impact on evolutionary outcomes. Our findings underscore the importance of precisely measuring the fitness costs of microbial social behaviors, and help explain the prevalence of the T6SS across Gram negative bacteria.

## Main Text

An important mechanism of intermicrobial antagonism is the Type Six Secretion System (T6SS). The T6SS is a modified phage tail spike that delivers a payload of deadly effector proteins into neighboring competitor bacterial species. Its widespread distribution (∼25% of Gram negative species [1]) can be understood through its outsized effect on microbial dynamics. The T6SS is both potent, being able to deliver toxins with up to a 99.99% fatality rate, and broad-acting, being able to affect Gram-negative bacteria, Gram-positive bacteria, as well as eukaryotes. The T6SS is utilized across a broad range of evolutionary niches, facilitating defense against competitors, as well as host invasion through the killing of commensal species [2].

The structure and function of the T6SS has been the subject of intense research since its discovery. With the exception of the baseplate and species-specific effector proteins, crystal structures have been determined for all other proteins comprising the canonical T6SS structure [3]. Studies have also explored the role of T6SS in other species where it has been repurposed for functions other than antagonism such as ion sequestration, biofilm formation, swarming, as well as various host-microbe interactions [4–7]. Given its hypothesized multi-megadalton size, complexity, and conservative developmental regulation, prior work has inferred that the T6SS must be metabolically costly to maintain and utilize [2, 4, 8, 9]. However, recent work attempting to measure the fitness cost of T6SS expression has found no detectable effect during a single 24 hour growth curve [10, 11]. Single-day growth curves have a relatively low sensitivity, however, as they can only measure competition outcomes over a limited number of generations, and are thus ineffective for measuring traits with a small impact on fitness [12]. To generate more precise estimates of the fitness costs of constitutive T6SS expression, we examined the outcome of competition between isogenic T6SS- and T6SS+ strains over 5 days (∼100 generations) of competition.

We created two strains of *Vibrio cholerae* [Supplemental 1], one with constitutive T6SS expression (T6SS+) and a T6SS(-) genotype that includes a transcriptional terminator placed in front of the T6SS operon. The difference in growth rate between these strains should therefore be due entirely to the metabolic cost of T6SS protein expression, T6SS assembly, and T6SS firing events. To differentiate these two strains during competition experiments, we also introduced a selectable antibiotic resistance marker for streptomycin (StrepR) into the lacZ gene of each strain (accounting for the effect of antibiotic resistance via marker-swap experiments) so that a blue-white screen can distinguish the two strains.

We measured the fitness costs of T6SS expression under two conditions: in liquid media, where cells do not directly interact [13], and on agar plates [Fig. 1a]. While both conditions include the growth-rate costs of T6SS expression, growth on solid media also introduces costs associated with friendly fire. Even though clonemates are resistant to the toxins delivered by the T6SS, sublethal toxicity and physical damage to cell walls and plasma membranes may incur an additional cost to T6SS expression when the population is growing as a spatially structured biofilm [14–16]. In both cases, we examined competition between T6SS+ and T6SS-strains over ∼100 generations. We ran each competition experiment twice, once with the T6SS+/StrepR and once with T6SS-/StrepR. We subtracted the results of the T6SS+/StrepR experiment from the T6SS-/StrepR experiment to isolate only the cost of T6SS.

**Figure 1.**
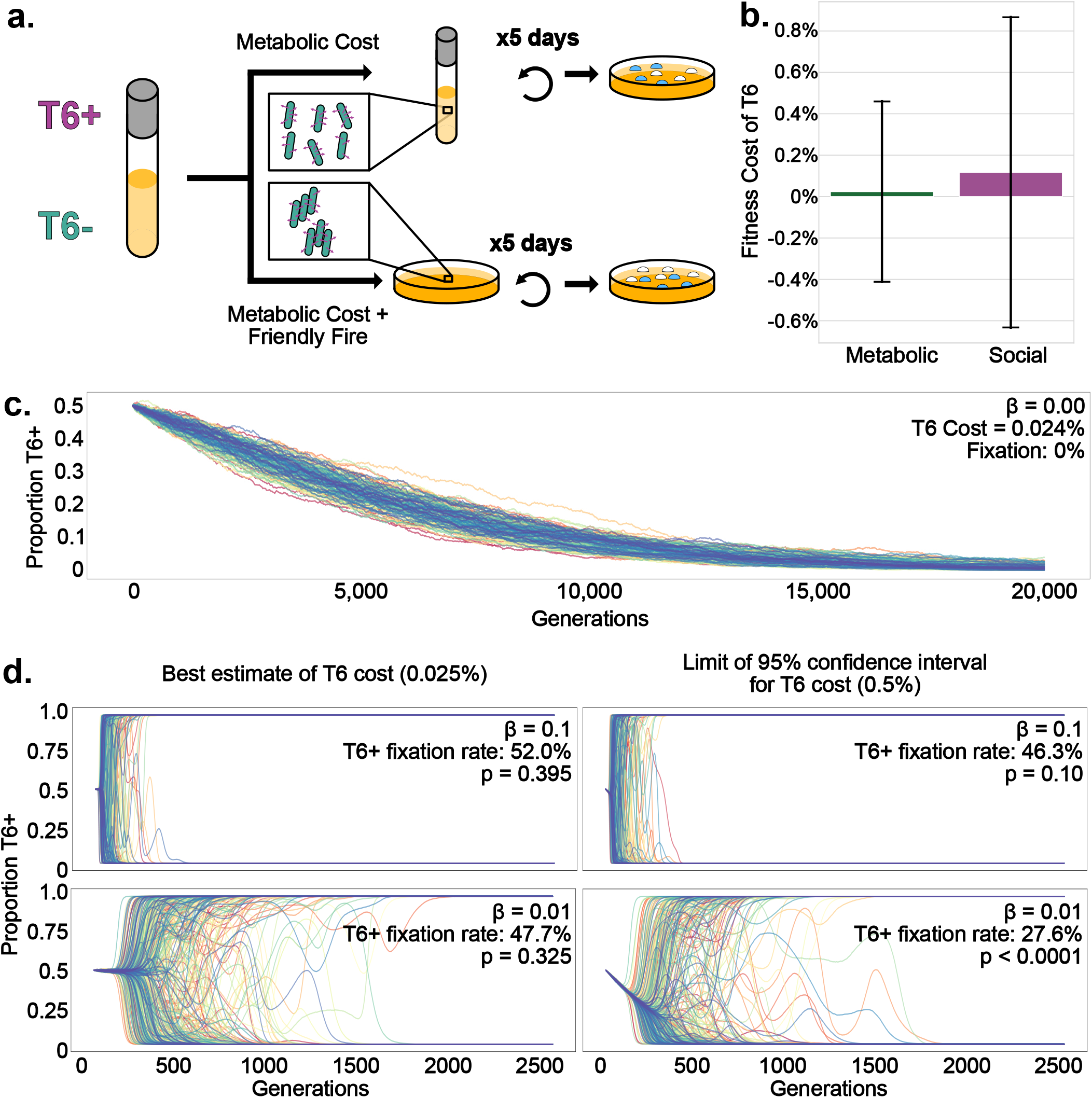
Constitutive expression of the T6SS is selectively neutral under most population genetic conditions. **a)** Experimental set-up for competitions. T6SS+ and T6SS-cells were initially mixed in an equal proportion. To measure the metabolic cost of constitutive T6SS expression, the mixture was incubated in liquid culture at 37°C with shaking and back-diluted daily, for five transfers. To measure any additional costs incurred by friendly fire, we performed competitions on LB agar at 37°C, and diluted every 8 hours for five days. The change in frequency of the T6SS+ strain was determined via a blue-white screen. **b)** In liquid media, the fitness cost of T6SS was 0.025% (error bars reflect the 95% CI; this value is not significantly different from 0). On solid media, the cost of T6SS was 0.11% (error bars reflect the 95% CI; not significantly different from 0). **c&d)** Simulations of 250 bacterial populations consisting of 1 million bacteria using a Wright-Fisher model. In the absence of clonal interference from beneficial mutations (*β*=0), all T6SS+ populations go extinct after >20,000 generations (C). Clonal interference (*β*>0) results in far more stochastic outcomes (d). The T6SS is non-neutral only when future beneficial mutations have a small effect (*β* = 0.01) and the cost of the T6SS is high (0.5%).

In liquid media, constitutive T6SS expression reduced fitness by 0.025 ± 0.5% (uncertainty represented as a 95% confidence interval), but this was not statistically distinguishable from zero (t=0.12, p=0.91, df = 16). On solid media, which includes costs of friendly fire in addition to the metabolic burden of T6SS expression, the effect was similarly small: 0.11 ± 0.75% (95% CI, t= 0.35 p= 0.73, df = 9) [Fig. 1b]. For comparison, the cost of the antibiotic resistance, also measured in this experiment, was 2.9% ± 0.2% (95% CI, t = 4.24, p = 0.0003, df = 16) [Sup. 1]. In our experiment, T6SS+ cells did not grow at a significantly different rate from T6SS-competitors. However, it is possible the true metabolic cost of T6SS was up to 0.5% (the 95% confidence interval of our liquid media experiments ranged from -0.5% to 0.5%).

We can contextualize these costs via simple mathematical models. Assuming the metabolic cost we measured of T6SS (0.025%) is true, a T6SS+ strain starting at 50% frequency in a population of a million bacteria would take more than 20,000 generations to go extinct [Fig. 1c]. Real microbial populations are more dynamic, however, as other mutations arise within these lineages and themselves become subject to selection. In asexual organisms like bacteria, this drives a phenomena known as clonal interference: competition between independent lineages in the population bearing different beneficial mutations [17]. Depending on key population genetic parameters (e.g., the rate and magnitude of beneficial mutations, population size, etc), mildly beneficial mutations that otherwise would have been fixed may be rendered effectively neutral due to clonal interference [17]. To explore the effects of clonal interference on selection against the T6SS, we created a stochastic Wright-Fisher model that examines the growth and competition of a population of T6SS+ and T6SS-bacteria subject to mutation and selection. We simulated a population of a million bacteria, starting with an even ratio of T6SS+ and T6SS-cells. We utilized the parameters from Good et al. [18]. Namely β, the magnitude of beneficial mutations, is 0.01; and m, the mutation rate per bacteria per generation, is 10^−5^. By varying the supply of beneficial mutations (β, larger values result in more clonal interference) as well as the cost of the T6SS, we examine the conditions under which the costs of T6SS are non-neutral.

Using our best estimate of the metabolic cost of constitutive T6SS expression (0.025%), we find that the T6SS has no effect on evolutionary outcomes at either β value simulated (Fig. 1d, left), suggesting that at the measured cost, even slow rates of adaptive evolution render the T6SS neutral. However, if we take the worst-case scenario, and assume that the true cost of the T6SS is 0.5% (the limit of our 95% confidence interval), then the cost of the T6SS is non-neutral when the supply of novel beneficial mutations is low (β = 0.01), but not when it is high (β = 0.1, Fig. 1d, right). Taken together, these results demonstrate that the T6SS is likely neutral under most population genetic conditions: the only time T6SS cost significantly alters the course of evolution in our simulations is either when T6SS has a statistically-improbable cost of expression and when the bacteria have a very low supply of novel beneficial mutations (as might be the case if they were near a fitness optima).

Our work demonstrates that the fitness costs associated with constitutive expression of the Type VI Secretion System in *V. cholerae* are low, and will often be imperceptible to natural selection, even when considering the additional costs incurred by friendly fire. In contrast, recent work from others has shown that T6SS was strongly selected against within animal models [10]. Together, these results suggest that while the expression of the T6SS is not metabolically costly, it may nonetheless be costly to express in certain environments (e.g., through triggering host immune response [19], or via emergent consequences of changed ecological dynamics).

This is notable as environmental *V. cholerae* isolates typically express far more T6SS than clinical isolates [20], which may reflect a difference in the cost of T6SS expression in these two environments. If our results hold for other bacterial species, then the low cost of T6SS, combined with the potential competitive benefits it provides, may help explain its widespread prevalence among Gram-negative bacteria.

## Acknowledgments

Christopher Zhang was funded by a GAANN fellowship from the Department of Education (P200A210046)

## Data Availability Statement

All data generated or analyzed during this study are included in this published article [and its supplementary information files].

## Supplementary Information

S1: .ods file containing spreadsheet of strains used, spreadsheet of measured results, values, and calculations

S2. .ipynb file containing code used for the model

S3: .pdf file containing methods

