## Supplemental Table 1 for "Constitutive expression of the Type VI secretion system carries no measurable fitness cost in *Vibrio cholerae*"

Strain list

| Strain Designation | Organism | Strain Background | Genotype | Markers | Notes |
|---|---|---|---|---|---|
| CZ0001 | V. cholerae El Tor | C6706str2 | Ptac-qstR | KanS, StrpR | Constitutive T6SS activity |
| CZ0002 | V. cholerae El Tor | C6706str2 | Ptac-qstR , T6SS large cluster promoter replaced with T4 terminator | KanS, StrpR | No measurable T6SS activity |
| CZ0003 | V. cholerae El Tor | C6706str2 | Ptac-qstR , Spec inserted into LacZ | KanS, StrpR, SpecR | Constitutive T6SS activity, white in a blue/white screen |
| CZ0004 | V. cholerae El Tor | C6706str2 | Ptac-qstR , T6SS large cluster promoter replaced with T4 terminator, Spec inserted into LacZ | KanS, StrpR, SpecR | No measurable T6SS activity, white in a blue/white screen |

### Metabolic Cost Competition Experiment

Quantity of bacteria counted on a plate

| | day 0 | | day 1 | | day 2 | | day 3 | | day 4 | | day 5 | | |
|---|---|---|---|---|---|---|---|---|---|---|---|---|---|
| | blue | whole | blue | whole | blue | whole | blue | whole | blue | whole | blue | whole | |
| Mixture 1 (CZ0001 and CZ0004) | 231 | 439 | 354 | 556 | 53 | 78 | 37 | 48 | 55 | 71 | 56 | 64 | |
| | 236 | 456 | 432 | 706 | 62 | 89 | 48 | 58 | 43 | 47 | 53 | 60 | |
| | 224 | 412 | 368 | 654 | 47 | 65 | 24 | 32 | 53 | 62 | 64 | 71 | |
| | 192 | 388 | 334 | 544 | 49 | 73 | 36 | 42 | 66 | 76 | 66 | 75 | |
| | 186 | 358 | 362 | 624 | 58 | 77 | 38 | 58 | 43 | 54 | 71 | 81 | |
| | 248 | 482 | 366 | 588 | 33 | 52 | 33 | 46 | 27 | 33 | 47 | 54 | |
| | 332 | 702 | 36 | 52 | 56 | 69 | 107 | 137 | 84 | 97 | 91 | 100 | |
| | 574 | 1104 | 27 | 38 | 38 | 53 | 86 | 102 | 51 | 62 | 99 | 106 | |
| | 372 | 758 | 23 | 39 | 51 | 67 | 69 | 92 | 75 | 90 | 80 | 95 | |
| Mixture 2 (CZ0002 and CZ0003) | 508 | 924 | 27 | 47 | 36 | 51 | 78 | 104 | 55 | 70 | 94 | 106 | |
| | 316 | 646 | 29 | 44 | 39 | 67 | 59 | 82 | 48 | 57 | 71 | 92 | |
| | 378 | 804 | 28 | 45 | 28 | 39 | 63 | 87 | 55 | 64 | 86 | 98 | |
| | 312 | 774 | contaminated | contaminated | 45 | 66 | 58 | 82 | 43 | 59 | 78 | 94 | |
| | 362 | 730 | 24 | 42 | 47 | 70 | 61 | 85 | 47 | 55 | 100 | 113 | |
| | 458 | 974 | 24 | 39 | 34 | 51 | 70 | 98 | 61 | 77 | 91 | 101 | |
| | 380 | 764 | 29 | 50 | 47 | 68 | 49 | 70 | 53 | 63 | 95 | 104 | |
| | 338 | 798 | 31 | 58 | 48 | 71 | 49 | 61 | 49 | 62 | 79 | 90 | |
| | 416 | 878 | 33 | 53 | 40 | 61 | 58 | 78 | 60 | 79 | 105 | 118 | |
| Dilution factor per day: | | 50 | | 5000000 | | 5000000 | | 5000000 | | 5000000 | | | 1.00E+06 |

TOTAL DILUTION FACTOR
3.125E+034

| | quantity of bacteria in solution (start of week) | | | quantity of bacteria in solution (end of week) | | | Malthusian – Blue (mBlue) | Malthusian – White (mWhite) | mBlue/mWhite |
|---|---|---|---|---|---|---|---|---|---|
| | blue | white | whole | blue | white | whole | | | |
| Mixture 1 | 23100 | 20800 | 43900 | 1.75E+036 | 2.5E+035 | 2E+036 | 14.6810182478422 | 14.3128121441953 | 1.02572563 |
| | 23600 | 22000 | 45600 | 1.65625E+36 | 2.1875E+035 | 1.875E+036 | 14.6657234735049 | 14.2748879723402 | 1.027379234 |
| | 22400 | 18800 | 41200 | 2E+036 | 2.1875E+035 | 2.21875E+36 | 14.7138788581005 | 14.3063250890446 | 1.028487663 |
| | 19200 | 19600 | 38800 | 2.0625E+036 | 2.8125E+035 | 2.34375E+36 | 14.7508633257993 | 14.3482534354207 | 1.028059854 |
| | 18600 | 17200 | 35800 | 2.21875E+36 | 3.125E+035 | 2.53125E+36 | 14.7718180924652 | 14.3954495750357 | 1.026144964 |
| | 24800 | 23400 | 48200 | 1.46875E+36 | 2.1875E+035 | 1.6875E+036 | 14.6317752229086 | 14.2625492585391 | 1.025887796 |
| | 33200 | 37000 | 70200 | 2.84375E+36 | 2.8125E+035 | 3.125E+036 | 14.7055763593196 | 14.2211757661392 | 1.034061923 |
| | 57400 | 53000 | 110400 | 3.09375E+36 | 2.1875E+035 | 3.3125E+036 | 14.6129295425626 | 14.0990380803014 | 1.03644869 |
| | 37200 | 38600 | 75800 | 2.5E+036 | 4.6875E+035 | 2.96875E+36 | 14.6570580078797 | 14.314874018127 | 1.023904087 |
| Mixture 2 | 50800 | 41600 | 92400 | 2.9375E+036 | 3.75E+035 | 3.3125E+036 | 14.6269941187379 | 14.2552757297049 | 1.026075847 |
| | 31600 | 33000 | 64600 | 2.21875E+36 | 6.5625E+035 | 2.875E+036 | 14.6658189844904 | 14.4135174084521 | 1.017504511 |
| | 37800 | 42600 | 80400 | 2.6875E+036 | 3.75E+035 | 3.0625E+036 | 14.6683220719264 | 14.2505249125033 | 1.029318019 |
| | 31200 | 46200 | 77400 | 2.4375E+036 | 5E+035 | 2.9375E+036 | 14.6871725795756 | 14.2918362180312 | 1.027661691 |
| | 36200 | 36800 | 73000 | 3.125E+036 | 4.0625E+035 | 3.53125E+36 | 14.707136646632 | 14.2958047356581 | 1.028772911 |
| | 45800 | 51600 | 97400 | 2.84375E+36 | 3.125E+035 | 3.15625E+036 | 14.6412295162801 | 14.1757271173021 | 1.032837991 |
| | 38000 | 38400 | 76400 | 2.96875E+36 | 2.8125E+035 | 3.25E+036 | 14.6871725795756 | 14.2137478567493 | 1.033307522 |
| | 33800 | 46000 | 79800 | 2.46875E+36 | 3.4375E+035 | 2.8125E+036 | 14.6737118431964 | 14.2177652084626 | 1.032068798 |
| | 41600 | 46200 | 87800 | 3.28125E+36 | 4.0625E+035 | 3.6875E+036 | 14.6890864697788 | 14.2503083450755 | 1.030790781 |

Mixture 1:

average
1.02845553778326

standard deviation
0.00413428062976378

count
9

Mixture 2:

average
1.02870423009332

standard deviation
0.00484461982782044

count
9

difference in averages
0.000248692

degrees of freedom
16

95% t statistic
2.120

pooled standard deviation
0.004503477

95% confidence range
0.004500674

bottom of ci
-0.00425198

top of ci
0.004749367

T-score
0.117144155

P = 0.906991

| Assuming T6SS does not have a cost: Calculations for the cost of the antibiotic: | |
|---|---|
| average antibiotic cost | 95% t statistic |
| 0.028579884 | 2.110 |
| stdev antibiotic cost | T-score |
| 0.004370888 | 4.24264068711929 |
| 95% ci | P = .000274 |
| 0.002173781 | |

| | day 0 | | day 1 | | day 2 | | day 3 | | day 4 | | day 5 | |
|---|---|---|---|---|---|---|---|---|---|---|---|---|
| | blue | whole | blue | whole | blue | whole | blue | whole | blue | whole | blue | whole |
| Mixture 1 (CZ0001 and CZ0004) | 370 | 658 | 594 | 846 | 746 | 1066 | 756 | 922 | 621 | 674 | 364 | 397 |
| | 302 | 600 | 654 | 968 | 418 | 582 | 312 | 368 | 206 | 251 | 115 | 119 |
| | 342 | 594 | 376 | 544 | 338 | 430 | | | 265 | 322 | 352 | 396 |
| | 334 | 602 | 474 | 780 | 384 | 552 | 538 | 658 | 627 | 776 | 427 | 483 |
| | 354 | 658 | 314 | 478 | 292 | 410 | 358 | 428 | 284 | 315 | 94 | 103 |
| | 232 | 456 | 580 | 776 | 380 | 496 | 240 | 276 | 148 | 172 | 122 | 138 |
| Mixture 2 (CZ0002 and CZ0003) | 274 | 534 | 532 | 804 | 183 | 280 | 160 | 214 | 322 | 374 | 97 | 107 |
| | 321 | 561 | 318 | 480 | 233 | 309 | 170 | 204 | 193 | 209 | 57 | 59 |
| | 279 | 552 | 354 | 578 | 204 | 270 | 173 | 197 | 441 | 473 | 422 | 431 |
| | 345 | 600 | 252 | 424 | 133 | 157 | 78 | 87 | 91 | 110 | 22 | 22 |
| | 264 | 360 | 320 | 496 | 168 | 233 | 134 | 164 | 398 | 443 | 178 | 189 |
| | 240 | 492 | 199 | 337 | 165 | 285 | 110 | 161 | 126 | 167 | 38 | 53 |

| Dilution Factor | 0 | 1.1 | 1.2 | 1.3 | 2.1 | 2.2 | 2.3 | 3.1 | 3.2 | 3.3 | 4.1 | 4.2 | 4.3 | 5.1 | 5.2 | 5.3 |
|---|---|---|---|---|---|---|---|---|---|---|---|---|---|---|---|---|
| | | 100000 | 10000 | 10000 | 100000 | 100000 | 10000 | 10000 | 500000 | 100000 | 100000 | 10000 | 300000 | 300000 | 300000 | 300000 |
| | | 100000 | 10000 | 10000 | 100000 | 100000 | 10000 | 10000 | 500000 | 10000 | 300000 | 10000 | 300000 | 300000 | 300000 | 300000 |

TOTAL DILUTION mix1
4.05E+072

TOTAL DILUTION mix2
1.215E+072

| | quantity of bacteria in solution (start of week) | | | quantity of bacteria in solution (end of week) | | | Malthusian – Blue (mBlue) | Malthusian – White (mWhite) | mBlue/mWhite |
|---|---|---|---|---|---|---|---|---|---|
| | blue | white | whole | blue | white | whole | | | |
| | 37000 | 28800 | 65800 | 1.4742E+075 | 1.3365E+074 | 1.60785E+75 | 32.51266485054 | 32.0826440944064 | 1.013403532 |
| | 30200 | 29800 | 60000 | 4.6575E+074 | 1.62E+073 | 4.8195E+074 | 32.322835700338 | 31.6537748530632 | 1.021136842 |
| | 34200 | 25200 | 59400 | 1.4256E+075 | 1.782E+074 | 1.6038E+075 | 32.5216987658474 | 32.1668867874217 | 1.011030349 |
| | 33400 | 26800 | 60200 | 1.72935E+075 | 2.268E+074 | 1.95615E+75 | 32.5650632821907 | 32.2028076201852 | 1.011249195 |
| | 35400 | 30400 | 65800 | 3.807E+074 | 3.645E+073 | 4.1715E+074 | 32.2507342519676 | 31.8119740533263 | 1.013792297 |
| Mixture 1 | 23200 | 22400 | 45600 | 4.941E+074 | 6.48E+073 | 5.589E+074 | 32.3873914127537 | 31.9881232122173 | 1.012481764 |
| | 27400 | 26000 | 53400 | 1.17855E+074 | 1.215E+073 | 1.30005E+74 | 32.0674566916982 | 31.6235208096708 | 1.014038155 |
| | 32100 | 24000 | 56100 | 6.9255E+073 | 2.43E+072 | 7.1685E+073 | 31.929462146216 | 31.3176417687187 | 1.019535966 |
| | 27900 | 27300 | 55200 | 5.1273E+074 | 1.0935E+073 | 5.23665E+74 | 32.3578988237181 | 31.5926906737054 | 1.02422105 |
| | 34500 | 25500 | 60000 | omitted due to no white cells | | | | | |
| | 26400 | 9600 | 36000 | 2.1627E+074 | 1.3365E+073 | 2.29635E+74 | 32.1963070067043 | 31.8418495335413 | 1.011131812 |
| Mixture 2 | 24000 | 25200 | 49200 | 4.617E+073 | 1.8225E+073 | 6.4395E+073 | 31.906529564552 | 31.7108643399933 | 1.00617029 |

Mixture 1:
average
1.013848997
standard deviation
0.003738647
count
6

Mixture 2:
average
1.015019454
standard deviation
0.007061386
count
5

difference in averages
0.001170458

degrees of freedom
9

95% t statistic
2.262

pooled standard deviation
0.005470528

95% confidence range
0.007493032

bottom of ci
-0.00632257

top of ci
0.008663489

T-score
0.353338414

P = 0.731998
