## Supplemental Code 1 for "Constitutive expression of the Type VI secretion system carries no measurable fitness cost in *Vibrio cholerae*"

```
import numpy as np
import random
rng = np.random.default_rng()
from tqdm import tqdm
```

```
def WF (N,A,G,S,M,P):
  '''
  Wright-Fisher Simulation:

  N = Total individuals
  A = Number of T6+
  G = Generations
  S = Cost of T6+
  M = Mutation rate per cell per generation
  P = Parameter of exponential distribution
  '''

  data = [] #stores all the generational data
  data.append([[[1], [N-A]], [[1-S], [A]]]) #first generation of bacteria


  for g in range (G):
    #using the last generation of bacterial counts to calculate the new generation
    original_fitness = [data[-1][0][0], data[-1][1][0]]  #fitness of the bacteria
    total_T6_minus_mutants = len(original_fitness[0])

    fitness = np.concatenate(original_fitness)
    numbers = np.concatenate((data[-1][0][1], data[-1][1][1]))

    #Generate new mutants
    WFweights = (fitness * numbers) / np.sum(fitness * numbers) #weights of each bacteria
    new_bacteria = rng.multinomial(N, WFweights) #next generation of bacteria

    if M == 0:  #if there are no mutants, save processor power on calculating mutants
      data.append([[[1], [new_bacteria[0]]], [[1-S], [new_bacteria[1]]]])
      continue

    #prune bacterial strains with 0 quantity of bacteria
    fitness = [fitness[:total_T6_minus_mutants], fitness[total_T6_minus_mutants:]]
    new_bacteria = [new_bacteria[:total_T6_minus_mutants], new_bacteria[total_T6_minus_mutants:]]
    WFweights = [WFweights[:total_T6_minus_mutants], WFweights[total_T6_minus_mutants:]]

    for k in range (2):
      WFweights[k] = [v for i, v in enumerate (WFweights[k]) if v != 0 and new_bacteria[k][i] != 0]
      fitness[k] = [v for i, v in enumerate (fitness[k]) if v != 0 and new_bacteria[k][i] != 0]
      new_bacteria[k] = [v for i, v in enumerate (new_bacteria[k]) if v != 0 and new_bacteria[k][i] != 0]

    #reset variables after updating the 0 quantity bacteria
    new_bacteria = np.concatenate(new_bacteria)
    total_T6_minus_mutants = len(fitness[0])
    WFweights = np.concatenate(WFweights)

    #generating new mutants
    mutant_count = rng.binomial(N, M) # #of new mutants
    new_mutants = rng.multinomial(mutant_count, WFweights) #identifying which strains were mutated
    subtracted = new_bacteria - new_mutants #subtracting those strains out
    subtracted = [subtracted[:total_T6_minus_mutants],subtracted[total_T6_minus_mutants:]]
    num_T6_minus = sum(new_mutants[:total_T6_minus_mutants])# total T6- bacteria

    #calculating fitness benefits of each new bacteria
    new_benefits = rng.exponential(P, mutant_count)
    T6_minus_benefits = (new_benefits[:num_T6_minus])
    T6_plus_benefits = (new_benefits[num_T6_minus:])

    new_mutants = [new_mutants[:total_T6_minus_mutants],new_mutants[total_T6_minus_mutants:]]

    #adding the fitness benefit of the mutation onto the mutational background
    background = []
    [background.extend((fitness[0][j]) * np.ones(new_mutants[0][j]))for j in range (len(fitness[0]))]
    T6_minus_benefits = background + T6_minus_benefits

    background = []
    [background.extend((fitness[1][j]) * np.ones(new_mutants[1][j]))for j in range (len(fitness[1]))]
    T6_plus_benefits = background + T6_plus_benefits

    #update the datatable
    updated_data = []

    for i, v in enumerate ((T6_minus_benefits,T6_plus_benefits)):
      updated_data.append([np.concatenate((fitness[i], v)) ,
                           np.concatenate((subtracted[i], np.ones(len(v))))])
    data.append(updated_data)


  return (data)
```

Colab paid products - Cancel contracts here
