## Supplemental Methods 1 for "Constitutive expression of the Type VI secretion system carries no measurable fitness cost in *Vibrio cholerae*"

**Bacterial strains and media:** Bacterial strains were grown aerobically at 37°C overnight in lysogeny broth (LB) (1% w/v tryptone (Teknova, Hollister CA, USA), 0.5% w/v yeast extract (Hardy Diagnostics, Santa Maria CA, USA), 1% w/v NaCl (VWR Life Sciences, Radnor PA, USA)) with constant shaking or on LB agar (1.5% w/v agar; Genesee Scientific, San Diego CA, USA) standing at 37°C. LB-X-gal was made by mixing in 40 µg/mL of X-gal (GoldBio, St. Louis MO, USA) to LB agar while it is liquid.

**Mutant Construction**: All V. cholerae mutant strains were made using the pKAS allelic exchange system described by Skorupski et al using pKAS32 [1]. All insertions and changes to phenotype were confirmed with PCR, antibiotic screening, and killing assays.

**Liquid LB Competition Experiment**: Overnight cultures of *V. cholerae* were normalized to an OD600 = 1 then mixed in a 1:1 ratio by volume. The mixture was then serially diluted and 100 µL of the $10^{-3}$ dilution was mixed into 5mL of LB and incubated overnight shaking at 37°C. 100 µL of the $10^{-5}$ dilution was plated onto an LB X-gal plate for quantification with blue-white screening. For the subsequent 4 days, the overnight mixture was serially diluted and 100 µL of the $10^{-5}$ dilution was mixed into 5mL of LB and the $10^{-6}$ dilution was plated onto an LB X-gal plate for quantification. Fitness was calculated by finding the ratio of Malthusian parameters as described in Lenski et al. [2]

**Solid Agar Competition Experiment**: Overnight cultures of *V. cholerae* were normalized to an OD600 = 1 then mixed in a 1:1 ratio by volume. The mixture was then serially diluted and 100 µL of the $10^{-5}$ dilution was plated onto LB X-gal as well as on LB. The X-gal plate was saved for quantification. The LB plate was incubated standing at 37°C. Every following 8 hours for 5 days, the LB plate was taken out of the standing incubator and all of the agar was scraped off of the plate and transferred into a 50 mL conical tube containing 10mL of LB. This conical tube was vortexed for 30 seconds and 300 µL of the supernatant was transferred into a 96 well plate and diluted such that the subsequent plate will have between 200 and 1000 cells on it. Every three time points were recorded via plating onto X-gal for quantification with blue-white screening. Fitness was calculated with the same calculation as the Liquid LB competition experiment.

**Statistics** The 95% confidence interval for both competition assays was calculated with a two sample t-test for the difference in two means. See supplemental table for details. The p value in Figure 1D was calculated by counting the quantity of populations that reached fixation or extinction after 5000 simulated generations and using a $\chi 2$ -test to test the null hypothesis that the T6SS+ strain will go to fixation 50% of the time.
